# A secreted proteomic footprint for stem cell pluripotency

**DOI:** 10.1101/2021.04.14.439804

**Authors:** Philip Lewis, Edina Silajzick, Helen Smith, Nicola Bates, Christopher A Smith, David Knight, Chris Denning, Daniel R Brison, Susan J Kimber

## Abstract

With a view to developing a much-needed non-invasive method for monitoring the healthy pluripotent state of human stem cells in culture, we undertook proteomic analysis of the spent medium from cultured embryonic (Man-13) and induced (Rebl.PAT) human pluripotent stem cells (hPSCs). Cells were grown in E8 medium to maintain pluripotency, and then transferred to FGF2 and TGFβ deficient media for 48 hours to replicate an early, undirected dissolution of pluripotency.

We identified a distinct proteomic footprint associated with early loss of pluripotency in both hPSC lines, and a strong correlation with changes in the transcriptome. We demonstrate that multiplexing of 4 E8- against 4 E6- enriched biomarkers provides 16 ratio abundances which are each robustly diagnostic for pluripotent state. These biomarkers were further confirmed by Western blotting which demonstrated consistent correlation with the pluripotent state across cell lines, and in response to recovery assays.

## Introduction

Although the clinical potential of regenerative medicine, including approaches using human pluripotent stem cells (hPSCs), remains unrivalled, major hurdles exist in translating promising protocols for treatments into scalable, reproducible commercial processes. In particular, maintaining hPSCs in their pluripotent state has proven challenging and labour intensive. Advances have been made in automating the culture and expansion of high quality hPSCs, such as by the development of the SelecT (TAP biosystems – Hertfordshire, UK), Freedom EVO (TECAN Trading AG, Switzerland) systems, or technologies developed by Tokyo Electron (Kyoto, Japan). However, a key issue in hPSC culture which remains to be overcome is the absence of a rapid, reproducible, non-invasive, and quantitative metric for cell pluripotency.

The most used techniques all have major drawbacks when it comes to their use in hPSC manufacture. For instance, using immunofluorescence for assessing the core pluripotency transcription factors and cell surface markers does not accurately pick-up initial loss of pluripotency in entire cultures, but rather relatively late loss of pluripotency in individual cells. Immunofluorescence is also imprecise, generally non-quantitative, cannot easily be integrated into the manufacturing process, and the tested cells are lost from culture in this terminal assay. Using flow cytometry for identifying loss of pluripotency markers is helpful but shows profound run variability and sacrifices cell product. Currently the most quantifiable metrics of pluripotency available are RNA-based assays on cells e.g. Pluri-test [1] and ScoreCard [2], or teratoma assays [3]. However, these metrics are not currently cost-effective, require sacrifice of pluripotent cell product, and take too long to provide results on ‘at-risk’ cultures, which have likely been permanently compromised by the time the results are available. The most rapid and cost-effective tool researchers currently have is analysis of cell morphology, however this is inherently subjective. While attempts have been made to automate and quantify the characteristics of hPSC morphology [4, 5], it remains a low-resolution way to measure pluripotency [6]. Thus, there are no completely fit for purpose quantitative assays which can pick up very early loss of pluripotency.

To address these limitations, we sought to identify secreted protein biomarkers in spent culture media within 48 hours of the onset of hPSC pluripotency loss. Although the secretome of hPSCs has been studied in the past [7–11], there is yet to be a comparative study aiming to identify proteins in media indicative of the pluripotent state, and its loss. It is well established that hPSC self-renewal relies on the signalling pathways down stream of FGF-2 and TGFβ family members [12–14]. After removal of these growth factors from standard hPSC culture medium the cells lose their pluripotency network, a prerequisite for lineage differentiation [15]. Therefore, we used the removal of FGF-2 and TGFβ from the medium to trigger loss of the pluripotent state, aiming to identify protein biomarkers which could be detected in spent medium while it is still possible to rescue cell product. Using LC-MS/MS of conditioned medium, we identified four E8- and four E6-enriched proteins in the secretome which are highly indicative of healthy pluripotent cultures or early pluripotency loss. Additionally, we demonstrate that by multiplexing proteins enriched in E8 against those enriched in E6, we can increase the scale and robustness of our detection of change in cell state between conditions, in a manner which could lead to the development of a highly sensitive assay for very early signs of pluripotency loss. Thus, we use the ratios of these proteins in E8 and E6 to indicate the pluripotent state or otherwise of the cultured cells. We further demonstrate the utility of these markers by returning E6 cultures to E8 medium, leading to a full recovery of pluripotency as determined by our markers, and confirmed by teratoma assays. Such indicators in the spent medium will be particularly useful in a scale up culture setting to prevent or reverse culture deterioration while avoiding cell sampling and loss of product and are amenable to simple detection assays.

## Results

### Incipient pluripotency loss after 48 hours of FGF2/TGFβ removal is at the limits of detection by flow cytometry and immunocytochemistry

We first confirmed the time scale of loss of conventional human pluripotency - associated markers. HESC (Man-13: [16]) and hiPSC (Rebl.PAT: [17]) lines (collectively referred to as hPSCs) were cultured in either pluripotency maintenance medium (E8 [18]), or identical medium lacking the pluripotency maintenance factors FGF2 and TGFβ1 (E6 [19]) for 48 hours, before analysis **(Fig 1.A)**. A previous study indicated that mRNA for lineage specific marker proteins is not definitively expressed until at least four days i.e., slower than loss of pluripotency-associated markers: - after as little as 48 hours following removal of FGF2/TGFβ[15]. Therefore, it was hypothesised that it should be possible to identify secretome changes associated with the loss of pluripotency at this early time point, enabling cultures to be identified at a point at which they can be rescued.

**Figure 1.**
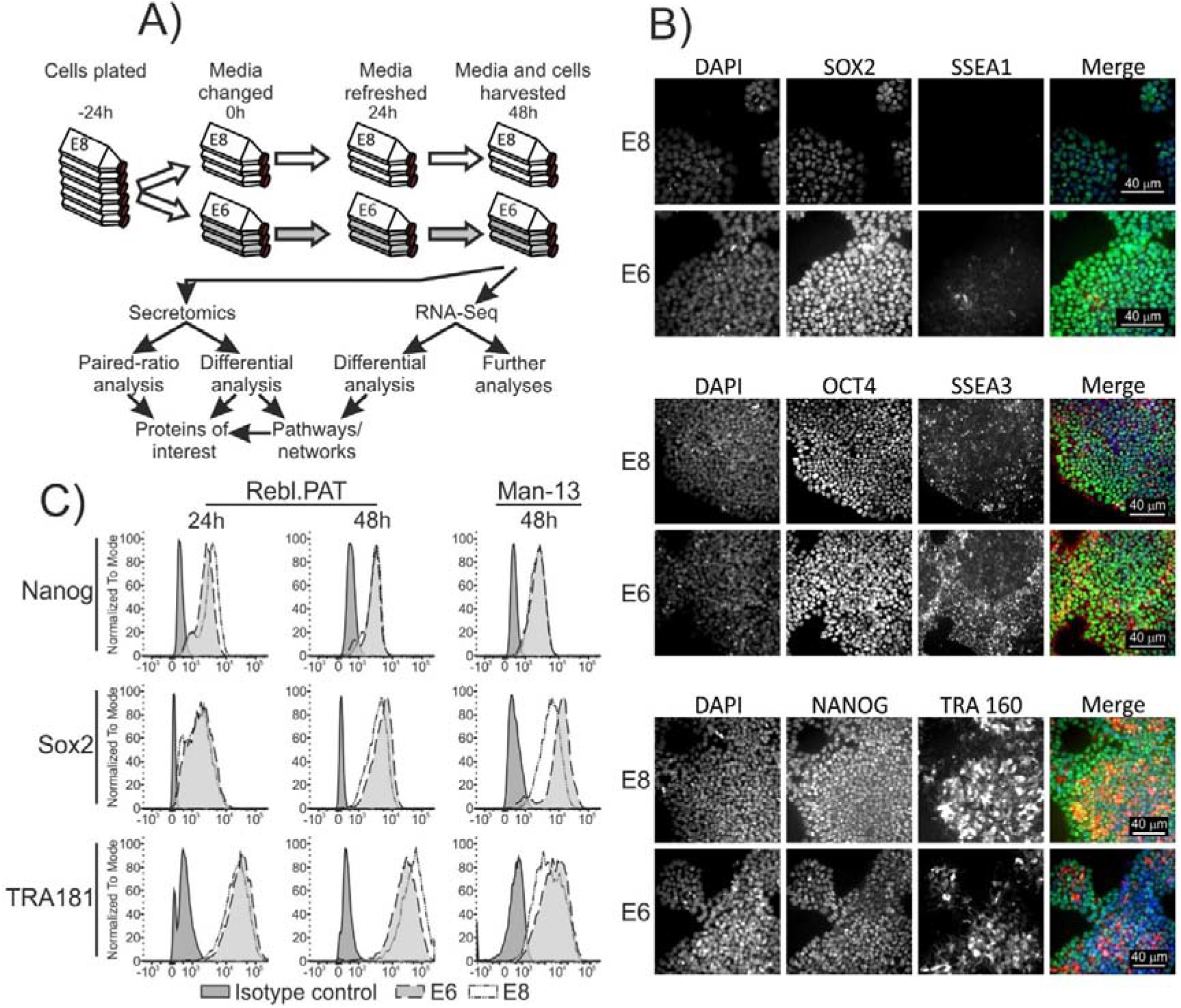
Experimental design for sample collection and data processing. A) Cells were cultured in E8 for at least two passages prior to the onset of the experiment. Cells were then plated in E8 medium containing Rock inhibitor onto Vitronectin-N coated flasks and were allowed to settle for 24 hours before onset of experimental conditions. Cells were washed in PBS, then half were cultured in E8, and half in E6, with full media replacement after 24 hours. After 48 hours of culture under experimental conditions the media was collected, spun, concentrated, and processed for LC-MS/MS. Cells were also collected and processed for quality control, and RNA-Seq analysis. B) Immunocytochemistry of Man-13 cells cultured in E8 (control) or after 48 hours of culture in E6. C) Flow cytometry of Rebl.PAT and Man-13 cells cultured in E8, or after 48 hours of culture in E6. For Rebl.PAT cells, extra flasks were cultured alongside experimental flasks, and these were used for observation of cell state at the 24-hour time-point (shown for Rebl.PAT).

After 48 hours of FGF2/TGFβ loss, MAN13 and Rebl.PAT cells in both E8 and E6 exhibited broadly comparable immunofluorescence (IF) and flow cytometry staining for pluripotency markers OCT4 and SSEA3, with a slight decrease in IF for NANOG and TRA160, and appearance of the early differentiation marker, SSEA1 in E6 **(Fig. 1B, C),** with NANOG and TRA1-81 very marginally decreased in E6 flow cytometry **(Fig. 1C)**. Only SOX2 was observed to be moderately increased after 48 hours in E6 **(Fig 1.B, C)** confirming transcript data **(Fig. S1)**. At 48hours, *OCT4* transcripts were slightly increased in Rebl.PAT as is common on initial differentiation. Additionally, no significant change in viability of cells between E8 and E6 was observed **(Fig. S2)**.

Although it is well established that feeder-free pluripotent stem cell maintenance requires FGF2 and TGFβ, without which cells lose pluripotency, our data indicate that current standard methods cannot unambiguously detect the incipient loss of pluripotency after 48h of growth factor withdrawal. Whilst the routine assays of flow cytometry Q-PCR and immunofluorescence can identify some subtle changes in pluripotency-associated markers at this time, the cells are only distinguishable when comparing these samples with high quality pluripotent controls and these changes would be almost impossible to identify in isolation as indicative of incipient pluripotency loss.

### Consistent, detectable changes in the secretome are observable after 48 hours of FGF2/TGFβ removal

Man-13 and Rebl.PAT cells were cultured in E8 medium for 3 passages before being plated for proteomic analysis. The protein was recovered from the collected medium and enriched for LC-MS/MS – in the workflow described in **Figure 1A** and **Materials and Methods.** In total we analysed 29 Rebl.PAT samples, and 24 Man-13 samples by label-free LC-MS/MS - the latter split into two proteomics runs of 12 samples each, termed Man-13 Run1 (M13R1) and Man-13 Run2 (M13R2). Differences in protein abundance between E8 and E6 samples were tested for statistical significance using Welch’s Two Sample t-tests (p) and adjusted for multiple comparisons using the Benjamini & Hochberg (q) method **(Fig 2A)**.

**Figure 2.**
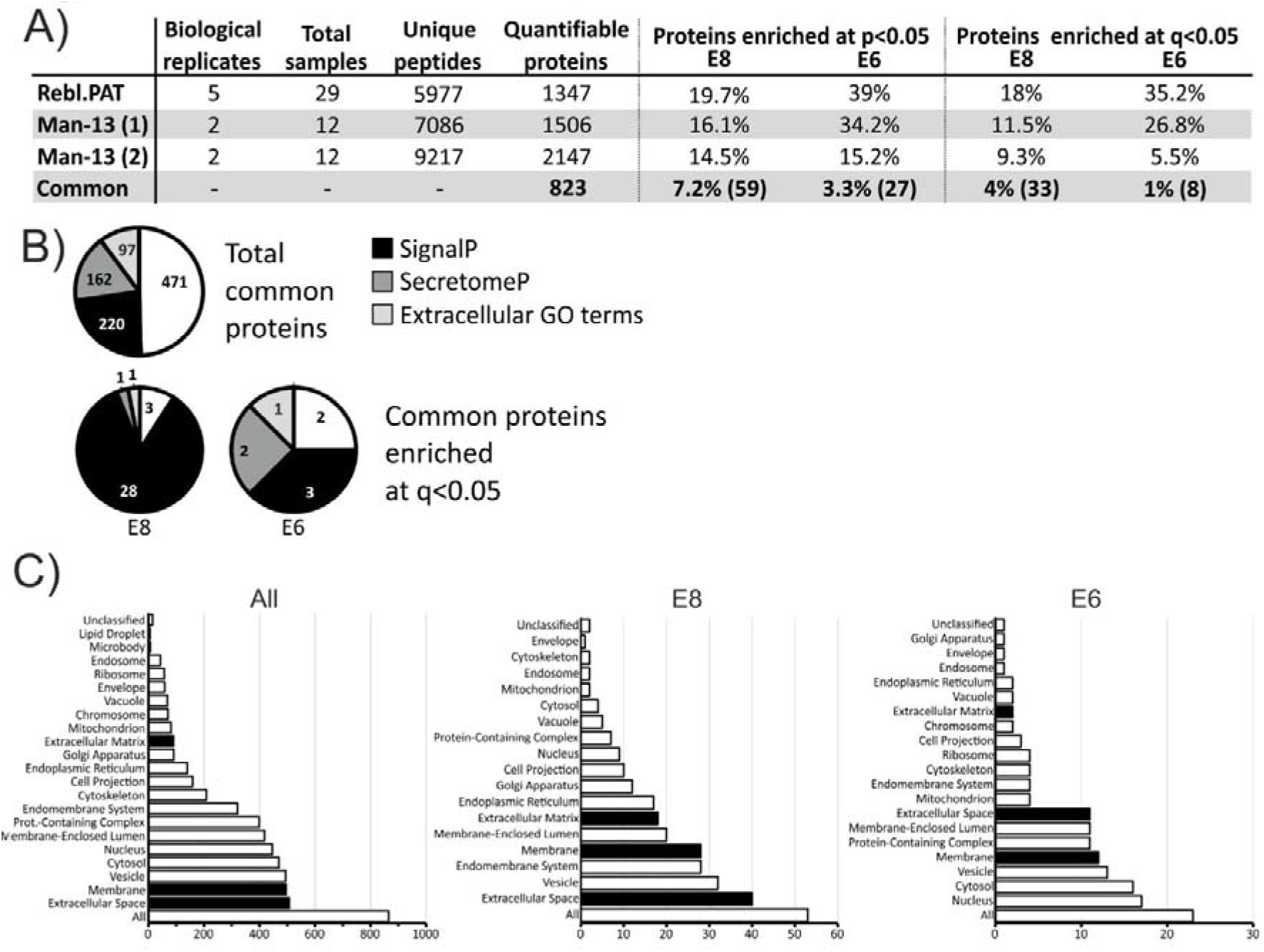
Proteomics summary. A) Summary of the proteomics experimental inputs and outputs. B) Pie charts demonstrating the proportion of proteins which are identified as secreted by SignalP 5.0, SecretomeP 2.0 or by extracellular Gene Ontology (GO) terms associated with their Uniprot accessions. Extracellular GO terms used for classification of proteins were extracellular matrix, extracellular space, extracellular vesicle, cell surface and extracellular region. All proteins used for this analysis had one or more unique peptides, and a confidence score of 30 or more in all three proteomics runs. C) GoSlim Cellular Component summary from WebGestalt of the popular GO terms for each of the categories described in C. Enrichment to p<0.05. Extracellular GO terms are highlighted with black bars.

Whilst more proteins were enriched (p<0.05) in E6 samples compared to E8, far fewer of these E6-enriched proteins were consistently changed across the three proteomics runs, and once adjusting for multiple comparisons **(Fig 2A)**. Moreover E8-enriched proteins (q<0.05) were more likely to be secreted, as identified by the presence of an amino-acid secretion tag (identification through SignalP [20]), an amino-acid sequence indicative of non-classical secretion (identification through SecretomeP [21]), or by annotation with the cellular component gene ontology (GO) tags; extracellular space, extracellular vesicle, cell surface, or extracellular region **(Fig. 2B)**.

Finally, the cellular component GO term membership of proteins identified in all three runs were compared between the total overlapping protein list, and the list of proteins enriched (p<0.05) in either condition across all replicates. This comparison demonstrated that extracellular GO terms were more common in E8-enriched proteins, whereas E6-enriched proteins were more likely to be associated with the terms Nucleus and Cytosol **(Fig. 2C)**. Taken together, this suggests that some of the proteomics changes we observed from E6 cultured cell-medium may be a mix both of pluripotency-loss cellular secretion and/or signalling proteins, and stochastic enrichment of intracellular proteins in the secretome which may be caused by a small increase in cell lysis or membrane leakage.

### Correlation between changes in protein and RNA abundance

We sought to confirm some of the proteomics changes we observed by looking at the transcriptomics of the same cell samples and correlating the fold change of the significantly changed (q<0.05) proteins and RNA **(Fig. 3)**. Firstly, we observed that proteins found to increase in E6 treated cells tended to also demonstrate an increase in transcript, and where disagreement in the direction of change between RNA and protein occurred, these proteins were often not identified as secreted. Changes in secreted protein and RNA abundance between E8 and E6 demonstrated a highly significant positive correlation (M13R1 R^2^ = 0.28, p<0.001; M13R2 R^2^ = 0.63, p<0.001; Rebl.PAT R^2^ = 0.59, p<0.001). Beyond further confirming the changes we observe in the proteome, these data also suggest the proteomic changes we observe indicate the onset of systemic changes in the pluripotent state of cells, rather than simply a transient effect restricted to immediate downstream signalling of FGF2/TGFβ1 or proteolytic action at the cell membrane,

**Figure 3.**
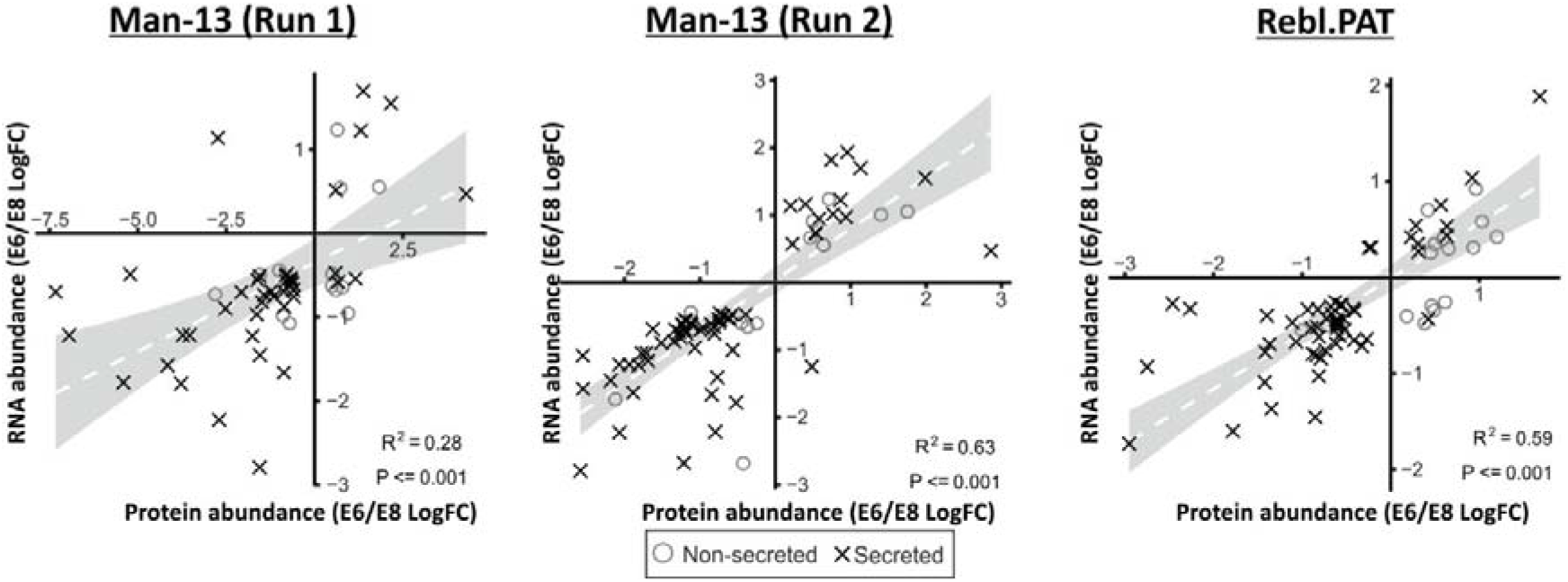
Correlation between E6 /E8 log fold-changes between mRNA and protein data. Protein data was paired to with RNA-Seq data using Entrez IDs retrieved from Uniprot and Bioconductor [78–80]. These data show that there is good agreement in the direction of change between proteomics data and the corresponding RNA-seq data. Only IDs which were q<0.05 significant by secretomic and RNA-Seq data were used for this comparison to avoid stochastic changes with little biological relevance. Man-13 samples processed for RNA-Seq were not split into two runs, as was the case in the proteomics dataset, as such the same RNA-Seq dataset was used to compare with both Man-13 proteomics runs.

### Selection of biomarker panel

As E8 and E6 media both contain high concentrations of transferrin, normalising the secreted proteome of stem cells against total protein content in the media is likely to introduce statistical uncertainty and error. Slight variabilities in cell density, starting viability, or growth rate could drastically change the relative contributions of cells and media to the total secretome, resulting in a changed normalised abundance of secreted proteins despite consistent secretion by cells.

Since there exists no protein secreted consistently enough to act as a normalisation standard, we sought to utilise the relative abundances of E6- and E8-enriched proteins. Since both proteins which are increased in abundance in E6, and proteins which are increased in abundance in E8 are responding to the same stimulus we expected that, regardless of changes in total protein abundance in the media, the relative abundances of these proteins are likely to remain consistent.

From the list of secreted proteins which demonstrate the largest, most consistent changes in RNA and protein abundance between conditions (**Table. 1**), we selected a panel of 4 proteins which increased when cells were cultured in E6 compared with E8 - Cochlin (COCH), FGF Receptor-1 (FGFR1), Follistatin (FST) and Olfactomedin-like 3 (OLFML3)-, and 4 proteins which increased when cells were cultured in E8 compared E6 - Chromogranin A (CHGA), Nidogen 1 (NID1), Neuronal Pentraxin-2 (NPTX2) and Semaphorin 3A (SEMA3A) (**Fig. 4**). It I worth noting that whilst FGFR1 is a membrane receptor protein, it is known to have an extracellular cleavage site [22], and all quantified FGFR1 peptides were extracellular of this site (**Fig. S3**).

**Table 1.**
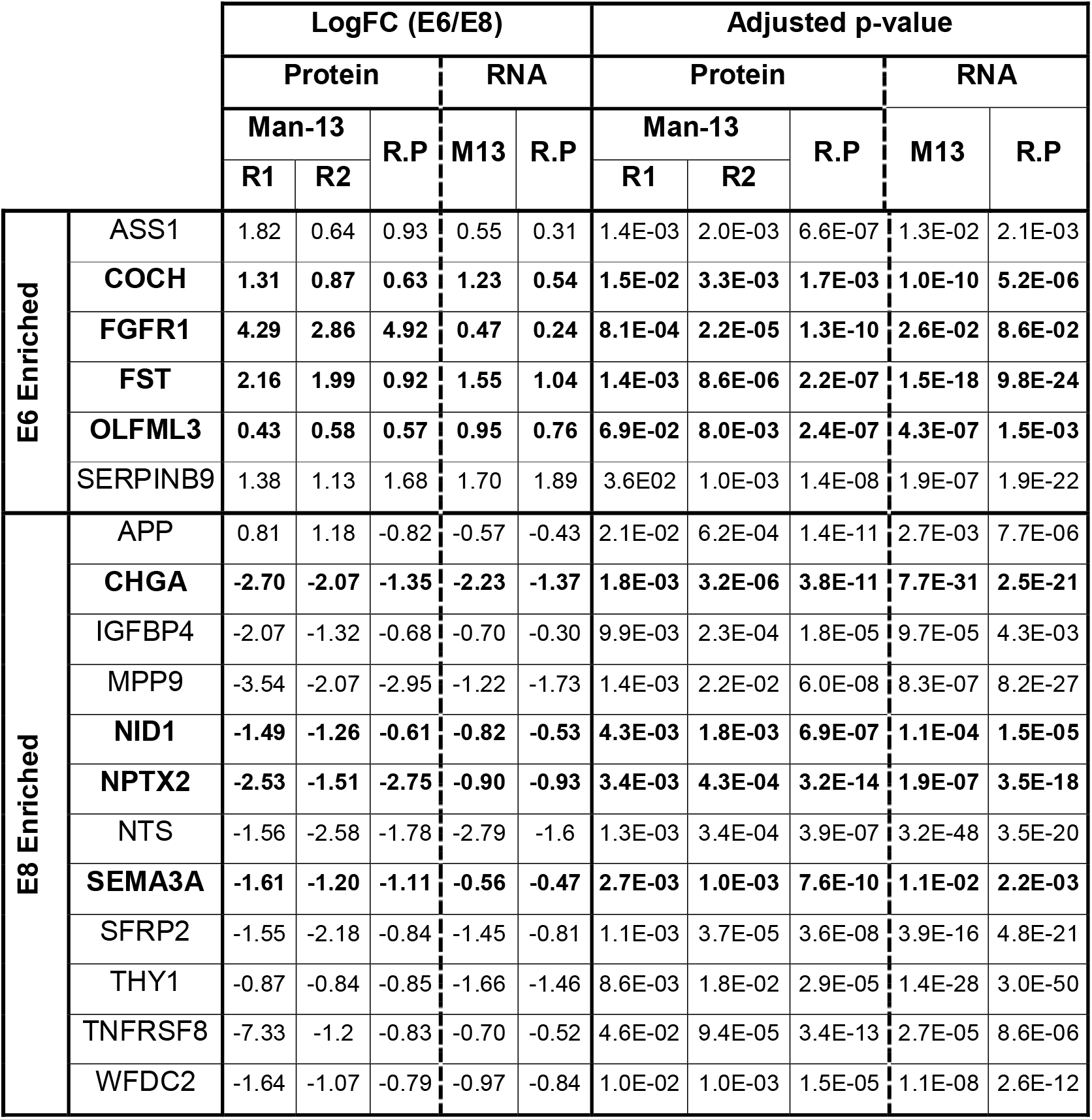
Proteomic and transcriptomics statistical summary for the top top candidate protein biomarkers. Proteins demonstrate a >1.5x increase in either E6 or E8 which passes a q<0.05 FDR cut-off in all three proteomics experiments. All proteins demonstrate evidence of secretion by the criteria listed in Materials and Methods and the majority have a matching q<0.05 change in the corresponding transcript. FGFR1 and OLFML3 are highlighted as FGFR1 does not have corroborating RNAseq data and OLFML3 misses the fold change cut-off in M13R1, however both are included due to the exceptional biological relevance of FGFR1, and the performance of OLFML3 by the paired-protein methodology demonstrated in Figure 4.

**Figure 4.**
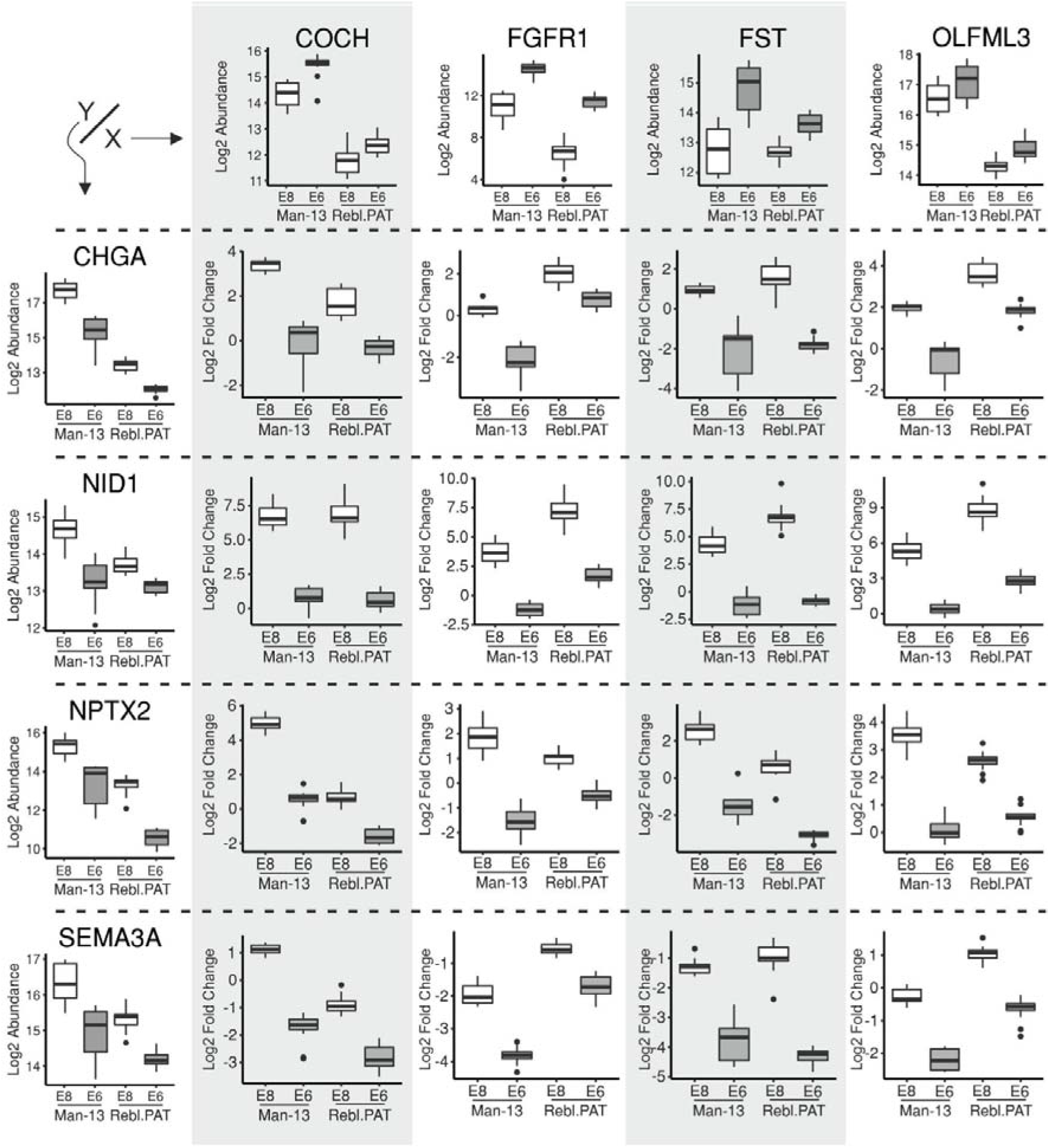
MS/MS individual abundance and relative abundance box plots for the marker proteins chosen for confirmation via Western blot. Individual abundances of single proteins (left column and top row) and relative abundance of protein pairs (intersection of the corresponding row/column) shown for the marker proteins selected for confirmation by Western blot based upon the paired relative abundances. Single protein box plot y axes show Log2-transformed individual abundance of individual markers. Relative abundance protein boxplots show Log2-transformed E6 marker individual abundance subtracted from the Log2-transformed E8 marker individual abundance. Box plot whiskers represent minimum and maximum values with the exception of outliers (shown as dots).

The 16 ratio pairs generated from comparing the 4 E8 against E6-enriched proteins are all diagnostic for incipient pluripotency loss across all samples in both cell lines (**Fig. 4**). It was noteworthy that data from both Man-13 LC-MS/MS runs were successfully pooled in this analysis, confirming that these marker ratios are consistent not only across both technical and biological replicates, but also different label-free LC-MS/MS experiments, giving confidence that the markers are robust to run to run variance.

### Confirmation of marker protein-pairs by Western blotting

Based upon the strength of these markers in the LC-MS/MS data, conditioned media samples (collected as described in **Figure 1A**) from the hESC lines, Man-1, Man-7, Man-13 and H9, and the hiPSC line, Rebl.PAT, were probed by Western blot for the relative abundances of these marker proteins (**Figs. 5 and S4**). Each protein-pair shown was probed for on the same membrane, and the abundance ratios calculated by densitometry measurements using ImageJ. These Western blots confirmed that the changes observed by LC-MS/MS are highly robust and discriminatory of pluripotent state across cells from different genetic backgrounds. The consistency of the ratio changes across multiple lines and between different combinations of proteins confirms this methodology as a discriminatory metric for incipient pluripotency loss across different hPSC lines.

**Figure 5.**
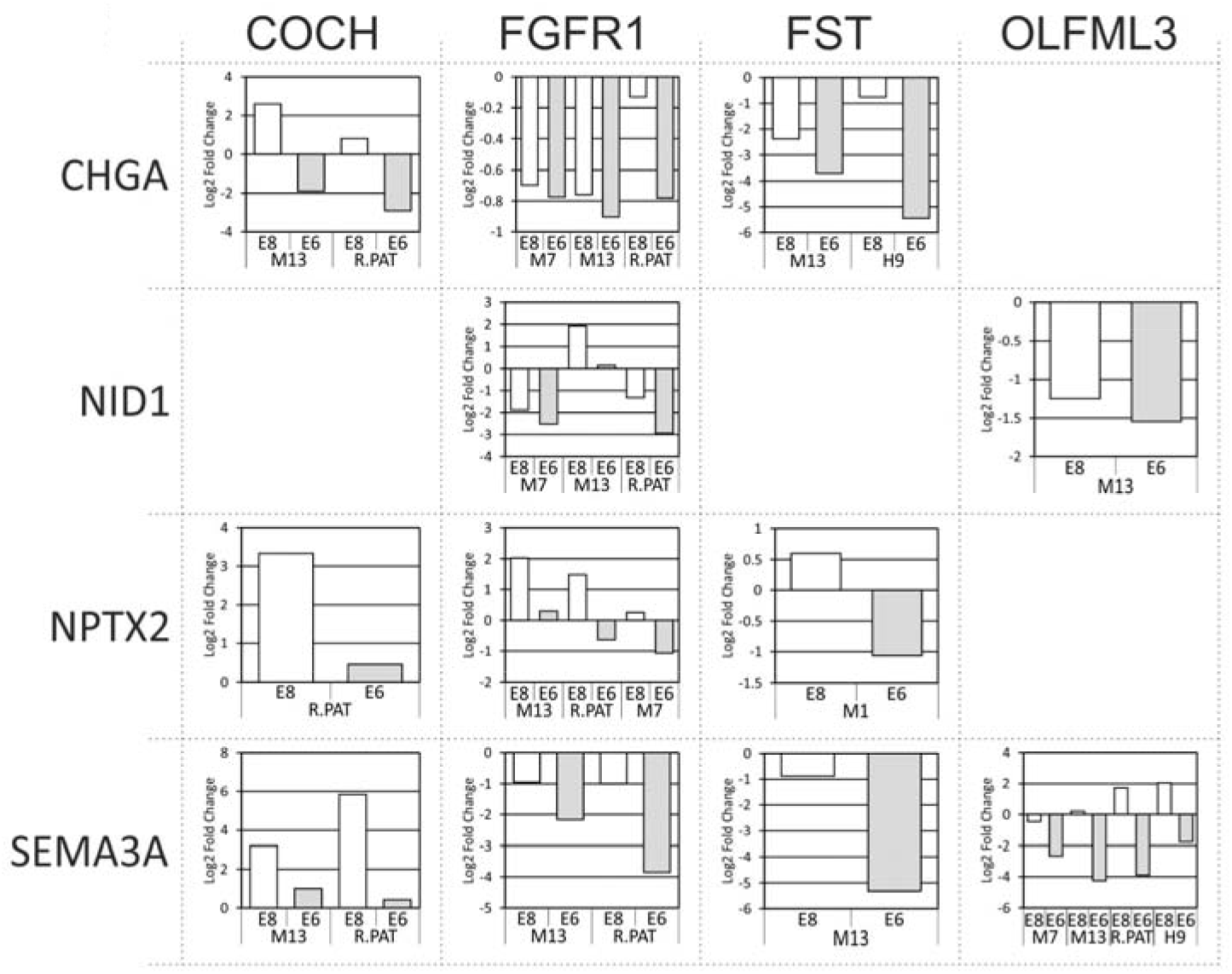
Confirmation of proteomics by Western blot on cell lines Man-1, Man-7, H9, MAN-13 and Rebl.PAT. Protein abundance was quantified by densitometry performed in ImageJ. The abundances are expressed as log2 ratios of E8-enriched proteins (left of the table) over E6-enriched proteins (top of the table). Equal quantities of concentrated conditioned media were used for each condition, and the same membrane probed with both antibodies for each ratio pair. Membranes were imaged using the LICOR Odyssey system. Images of the Western blots are available in Figure S2. B, C.

### Recovery of protein-pair ratio abundances after 48 hours in E6 medium

An important component of an assay for loss of pluripotency is the ability to detect a degradation of culture quality while the cells are recoverable. We therefore repeated the experiments indicated above but this time replaced some cells in E8 medium for 5-7 days after the 2 days in E6. Using Man-1 cells we demonstrated that the ratio between two pairs of the protein biomarkers selected from our analysis (OLFML3-NID1 and FST-NPTX2) returned to their pluripotent levels after 7 days of E8 recovery (**Fig. 6 A, B**). Moreover, cells cultured for 48 hours in E6 and recovered for 7 days in E8 media can still form teratomas (**Fig. S5**) demonstrating differentiation to tissues of all three germ layers. This suggests not only that the pluripotency of these cells can be fully recovered after 48h in E6, but also that our marker proteins can robustly track this recovery. That the changes we observe are not simply molecular markers of the immediate loss of growth factor is supported by the fact that the proteins take several days to return to their baseline, rather than returning immediately after the FGF2 and TGFβ1 are restored. Thus, at the time early loss of pluripotency is recognised by these early marker changes, cultures are still plastic and can recover.

**Figure 6.**
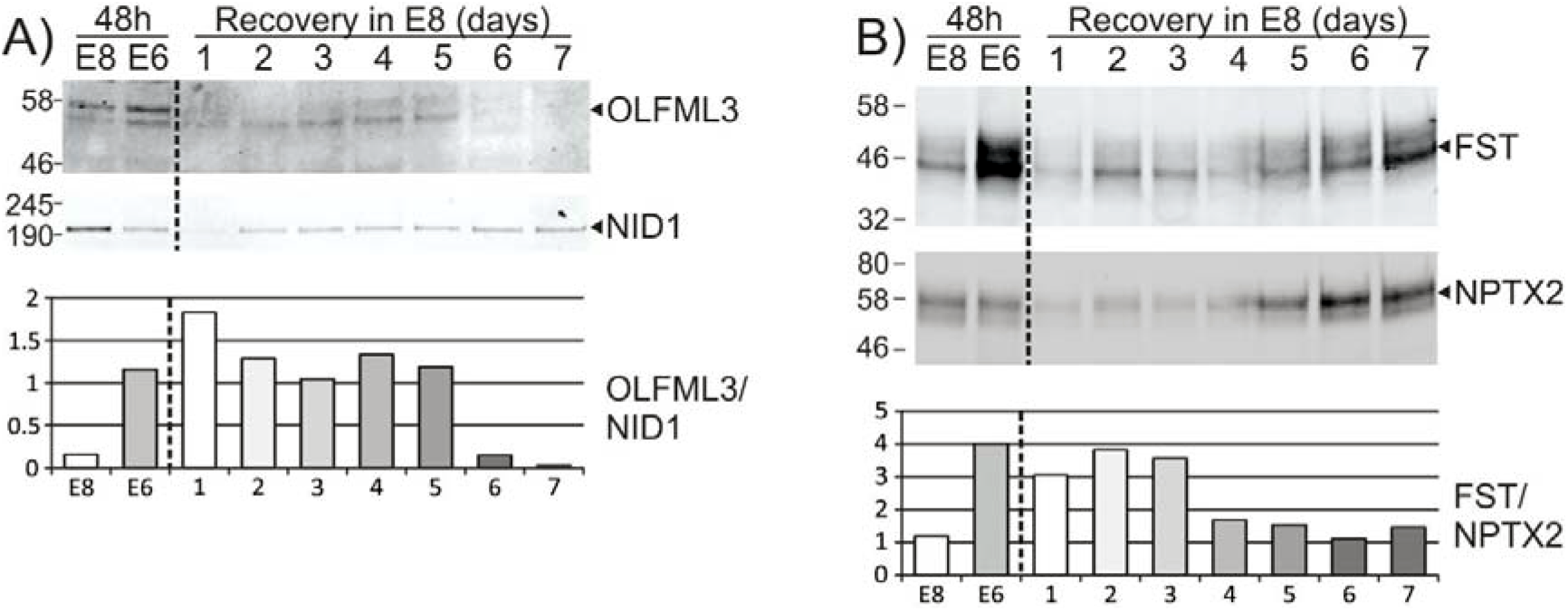
Re-establishment of relative marker abundances after a 7-day rescue. Man-1 cells were cultured as described in (Fig. 1.A), but after media collection at 48 hours, E6 cultured cells were passaged and returned to E8 for 7 days with media collection every 24 hours. Each Western blot membrane was probed with a pair of antibodies **(A)** OLFML3 & NID1 and **(B)** FST & NPTX2. Quantification shows the Log2-transformed E6 marker individual abundance subtracted from the Log2-transformed E8 marker individual abundance, with abundances calculated from densitometry. Densitometry was performed in ImageJ. Membranes were imaged using the LICOR Odyssey system. Densitometry was calculated in ImageJ. Membranes were imaged using the LICOR Odyssey system.

### Assessment in TeSR1 and mesodermal differentiation medium

In order to further validate that these secretome markers could be predictive with cells cultured in other commonly used media we grew Man13 in the pluripotent stem cell culture medium, TesR1, (**Fig.S6**) and chondrogenic differentiation medium [23]. TesR1 medium has high HSA which interferes with Western blotting and so this had to be removed before running the gel. Three different samples of TESR1 were concentrated to 50μg, 100μg and 200μg in total (after HSA removal) and were assessed by western blotting NPX2 was observed in all lanes in the TeSR1 medium after hESC culture (**Fig S6B**). A clear Follistatin band was detected strongly in the E6 medium using only 20μg protein. Here it is stronger than with 50μg protein from the TESR1 pluripotency medium. This confirms the enhancement of Follistatin on transfer of stem cells to a pro-differentiation medium (**Fig. S6A**). Similarly, when cells were transferred to a chondrogenic differentiation medium for 48h after culture in TeSR1 there was an increase in Follistatin and decrease of NPTX2 (**Fig. S6B**). NID1 was detected in TeSR1 after hESC culture but only at 200ug total protein and it is possible that this may be partly removed with the HSA.

## Discussion

We first investigated changes in commonly employed pluripotency associated proteins after only 48 hours following FGF2/TGFβ removal from the stem cell culture medium, confirming that neither immunofluorescence nor flow cytometry for these markers could confidently detect pluripotency loss at this time. Although it is a pluripotency-associated marker, SOX2 increased slightly. This is to be expected as SOX2 expression has previously been found to increase upon SMAD2/3 inhibition and FGF2 deprivation [24]. Moreover, increases in SOX2 in hPSCs can initiate differentiation [25, 26] and SOX2 has an important role in specification and differentiation of PSCs towards the neurectoderm lineage [27–29].

In this study we identified a series of proteins which function as secreted, early marker proteins for incipient loss of a pluripotent stem cell culture. To examine if the identified protein changes indicated a change in cell synthetic activity, we also investigated gene transcription and found that the proteome changes were indeed also generally reflected in changes in the respective gene transcripts. From this list of marker proteins (Table. 1) a shortlist of eight proteins was identified for further analysis by Western blotting.

### Biological relevance of marker proteins

Although the cellular proteomes of both hESC and hiPSC lines have been investigated by a variety of mass spectrometry techniques in the past [7–10] the protein secretome of these cells has been little studied [11]. Using a combination of LC-MS/MS and bioinformatic analysis with confirmation by Western blotting, we identified several markers in the secretome indicative of healthy pluripotent stem cells and early incipient pluripotency loss. Importantly these secretome markers were subsequently validated independently in hESC lines other than the lines Rebl.PAT and Man13 selected for the LC/MS interrogation. Moreover, we have also shown that cells returned to E8 after 48h in E6 show recovery and exhibit secretome marker proteins correlating with the pluripotent state again.

The transmembrane FGF2 receptor FGFR1 was one of the most highly represented secreted markers identified to increase upon growth factor withdrawal. There are several possibilities to explain the increased incidence of FGFR1 in the conditioned E6 medium. Firstly, FGFR1 is known to be released into the extracellular space by cleavage in its membrane-adjacent extracellular domain by MMP2 [22]. As FGFR1 proteins are endocytosed by the cell upon substrate binding and degraded in lysosomes [30–32], it is likely that decreased FGF2 abundance will result in an increased surface abundance of FGFR1, and a corresponding increase in the amount of FGFR1 which is cleaved by surface metalloproteinases. Indeed, a majority of the FGFR1 peptides identified as significantly increased in E6 in our proteomics study were N-terminal to the proposed MMP2 cleavage site. As it has previously been reported that a secreted form of the extracellular region of FGFR1 inhibits FGF2 activity [33] it could be speculated that the FGFR1 accumulation in FGF2 deficient media may further inhibit residual FGF2 signalling.

Indeed, the biological functions of the E6-enriched protein biomarkers are consistent with a hypothesis that cells triggered towards differentiation release factors into the medium that bind and modulate growth factors, sequestering pluripotency promoting factors such as FGF2 and TGFβ 1 to reduce their local availability. This will generate a feed-forward amplification of the original pluripotency dissolution signal which again highlights the need for early warning of incipient pluripotency loss. Follistatin is a well-known stem cell differentiation factor, and TGFβ inhibitor, functioning through the binding and neutralisation of TGFβ superfamily members [14]. Similarly, OLFML3, also enriched in E6, has been shown to bind to and stabilise BMP4, enhancing SMAD1/5/8 signalling in endothelial cells [34]. As BMP4 initiates differentiation of human embryonic stem cells to trophoblast cells in the absence of FGF [35–37] and to mesoderm and chondrocytes [38, 39], it could be concluded that after 48 hours of FGF2/TGFβ loss, these cells are secreting factors to promote and commit to the differentiation of nearby cells [40].

SEMA3A, a neuronal signalling protein involved in axon guidance, is highly enriched in the E8 condition by both secretomics and RNAseq, whilst its transmembrane receptor, NRP1 is highly enriched (q<0.01) in the E6 transcriptome [41–43]. This is significant as NRP1 also binds to both FGF2 and TGFβ and modulates their signalling. In HUVEC cells, increased NRP1 expression was demonstrated to suppress Smad2/3 activation upon TGFβ stimulation, and expression of SMAD target genes in response to TGFβ was increased in NRP1 deficient HUVECs [44], a relationship which has been replicated *in vivo* in murine endothelial cells [45]. NRP1 is a highly promiscuous receptor, and ligands compete for NRP1 binding such that competitive inhibition between ligands is common [46]. It has been demonstrated that ligand-competition for NRP1 exists between SEMA3A and VEGFA [46], and between VEGFA and TGFβ [47]. It could be tentatively suggested therefore that SEMA3A may also directly compete with TGFβ for NRP1 binding in a manner which prevents TGFβ sequestration and negative modulation by NRP1. In this way it is possible that it is protective for TGFβ signalling. However confirmation of this will require further research.

CHGA is a precursor protein for several neuroendocrine signalling proteins. Peptides from across the whole CHGA molecule were almost universally identified as being highly enriched in E8 conditioned medium over those in E6 conditioned medium. However, the association with the pluripotent state is unclear. Generally speaking, CHGA drives formation and release of secretory granules [48, 49], and its decreased abundance in stem cells during pluripotency dissolution could reflect a reduction in overall secretion. This would be consistent with the greater number of secreted proteins identified in E8 conditioned medium compared with E6 though further data is needed to confirm this association.

NID1 is a secreted glycoprotein with two principal protein-binding domains (G2 and G3), separated by a flexible chain [50]. By domain-specific binding of different components of the basement membrane, NID1 stabilises the basement membrane, cross-linking its multiple components [51, 52]. NID1 is abundant in locations where there is a requirement for additional resilience against mechanical stress, and conversely, less abundant where more flexibility is required and tissue disposition is not fixed [53] for instance being actively degraded during basement membrane disassembly [54]. It has been demonstrated by proteomic analysis to be expressed at significant amounts by hESCs [11, 55]. In our current data we observed a dramatic reduction in the abundance of NID1 in E6 media compared with E8. As NID1 has a stabilising effect on the basement membrane, it is likely that its abundance is decreased prior to the epithelial-mesenchymal transition inherent in the progression from a pluripotent stem cells to an early progenitor. This will involve the disassembly of several extracellular matrix complexes: in ovarian cancer cells NID1 plays a key role in this EMT transition [56].

Although COCH is known to be secreted [57], relatively little is known about its function beyond its role in the extracellular matrix of the inner ear [58]. In murine ES cells COCH has been reported to be expressed in response to BMP4 signalling and this was suggested to support self-renewal [59]. In our study COCH was enhanced in E6 medium possibly reflecting loss of FGF which is known to suppress BMP signalling in hESCs [12]. The acknowledged molecular difference in the pluripotent ESC state between human and murine reflect an earlier naïve murine ESC phenotype and later primed human ESC phenotype [60] and are supported by the wealth of data on the differences in signalling required for murine and human stem cell maintenance [12, 13, 61, 62]. Thus, BMP signalling has different effects in hESCs and murine ESCs, being active in stem cell maintenance in the latter [36, 62]. Since BMP signalling is strongly associated with differentiation to mesodermal and trophectodermal lineages in hESCs [35, 38, 63–65], this would be in keeping with a correlation between targets such as COCH down stream of BMP and the dissolution of pluripotency precipitating incipient lineage differentiation.

There is very little information on the function of NPTX2 outside of it’s binding to α-amino-3-hydroxy-5-methyl-4-isoxazolepropionic acid (AMPA) receptor subunit Glutamate receptor 4 (GRIA4) in neuronal tissues [66]. It is certainly noteworthy that the expression of *GRIA4* is also much higher in E8 than E6 (Supplementary RNAseq data) as for NPTX2, suggesting some as-yet unidentified function for this ligand-receptor pair in hPSCs

### Development of paired-ratio method

A caveat of these analyses, common to all secretome studies, is the lack of a suitable normalisation standard against which to compare proteins to robustly identify changes between conditions. Typically, ‘omics’ studies normalise to total protein or mRNA, or to house-keeping genes or proteins such as GAPDH or β-actin. These controls allow for reliable, reproducible comparisons across conditions or experiments. When analysing the secreted complement of the cell proteome, however, no such house-keeping protein exists, and the overall protein abundance between conditions is highly susceptible to variability due to the presence and concentration of medium proteins, the density of cells, differing cell viability, cytoplasmic leakage, and a host of other factors. In this study marker proteins were identified which could be multiplexed against each other to provide ratios which are much more discriminatory between the experimental conditions. Regardless of other changes in the cell secretome, the relative abundances of these proteins should remain relatively constant within an individual genetic background and cell state. As such, once an individual cell-line’s baseline ratios are identified under control conditions, it is expected that these ratios would provide a rapid, quantitative warning of imminent pluripotency loss in cultured hPSCs, above and beyond any current method in terms of sensitivity, affordability, and rapidity of timescale.

In summary we have shown that there are several proteins in the secretome of hPSCs that are rapidly decreased or increased during dissolution of pluripotency. By employing a matrix of four of each such species, robust ratios of proteins could be generated which were indicative of stable pluripotency or its incipient loss. Additionally, further such ratios of other proteins were identified by MS/MS and will make excellent targets for follow-up analyses. Such proteins will be invaluable as markers allowing intermittent or continuous culture monitoring during the scale up and manufacture of hPSCs for cell therapy or for their use in pharmaceutical drug development and toxicity testing. However, given cost effective means of monitoring such markers they will be equally valuable in the research laboratory, allowing rapid evaluation of culture status without loss of cell product in either context.

## Methods

### Cell culture

Cells were maintained in Essential 8 (E8) medium (Life Technologies) on Vitronectin-N (Life Technologies) and passaged using TrypLE dissociation reagent (Life Technologies). Prior to sample collection for secretome analysis, cells were plated in E8 medium, supplemented for 24 hours with 10μM ROCK inhibitor, at a density of 20,000 (Rebl.PAT) or 25,000 (Man-13) cells per cm^2^ in 75cm^2^ culture flasks (Corning). Twenty-four hours after plating (experimental time 0), cells were either fed with fresh E8 medium, or transferred to Essential 6 (E6) medium (Life Technologies). Cells were fed at 48 and 72 hours (experimental time 24h and 48h respectively) after plating and at these time-points medium was collected, spun at 1,000 x g at −9°C for 5 minutes then aliquoted into lo-bind 2ml centrifuge tubes (Eppendorf) and snap frozen in liquid nitrogen. At 72 hours after plating (experimental time 48h), cells were dissociated with TrypLE, spun at 1,000 x g in PBS for 5 minutes, the supernatant removed, and the pellet snap frozen in liquid nitrogen. Viability assays were conducted at medium harvest using Apotox-Glo assay (Promega) [67] or with the NucleoCounter NC-200 and Via-1 cassettes (Chemometec) according to manufacturers’ instructions.

A subset of the cells at this time-point were also collected and processed for RNA-Seq analysis. Five biological replicates were conducted for Rebl.PAT, each of which was composed of three technical replicates, and all samples were analysed by LC-MS/MS in a single run. To ensure quantitative reproducibility of the marker proteins identified across multiple analyses, four Man-13 biological replicates were collected, and these were analysed in two independent LC-MS/MS runs consisting of two biological replicates each. Some Man13 cells were also cultured in TeSR1 medium (Stem Cell Technologies) and then transferred to E6, or chondrogenic medium (DMEM:F12, 2mM L-glutamine, 1% (vol/vol) ITS, 1% (vol/vol) nonessential amino acids, 2% (vol/vol) B27, 90μM β-mercaptoethanol (all Life Technologies) + 20ng/ml BMP2 (R&D systems) 10ng/ml ActivinA (Peprotech, 20ng/ml FGF2 (Peprotech), 3μM CHIR99021 (Tocris), 100μM Pik90 (Selleckchem), modified from Wang et al 2019 [23].

An important component of an assay for decreased PSC culture quality is the ability to detect incipient pluripotency loss while the cells are still recoverable. The experiments above were repeated but after 2 days in E6, cells were restored to E8 medium for 7 days to assess whether full pluripotency could be recovered at the time that protein ratios detected early changes. Medium was refreshed every 24hrs with samples of conditioned medium being taken for analysis by Western Blot every 48 hours.

### Immunochemical staining of pluripotency markers

Cells were seeded at a density of 20,000 (Rebl.PAT) or 25,000 (Man-13) cells per cm^2^ in a 24 well plate and treated identically to samples prepared for proteomic and RNA-Seq analysis (**Fig. 1A**). Cell surface markers and transcription factors characteristic of pluripotent hESCs were detected using immunofluorescence. The cells were fixed with 4% paraformaldehyde and incubated with antibodies against stage specific embryonic antigens SSEA-4, SSEA-1 (R&D Systems), TRA-1–60, TRA-1-81 (Abcam) and transcription factors SOX2, NANOG (Cell signalling Technologies), and OCT-4 (BD Biosciences) at 4°C overnight. Secondary antibodies (Life Technologies), specific for the species and isotype of the primary antibody, conjugated to Alexafluors 488 or 594 were used for detection using a BX51 microscope (Olympus, Hertfordshire, UK) equipped with a Q-Imaging camera (Micro Imaging Applications Group, Inc, Buckinghamshire, UK). Image processing was done with the aid of Q-Capture Pro software package (Micro Imaging Applications Group, Inc).

### Gene expression analysis

Total RNA was extracted using mirVana™ miRNA Isolation Kit (Life Technologies), reverse transcribed using M-MLV reverse transcriptase (Promega) and candidate genes expression (normalised to GAPDH) assessed using SYBR Green PCR Master Mix (Applied Biosystems) with an ABI PRISM 7500 Real Time System (Applied Biosystems). Primers for rtPCR were as follows: GAPDH -FW (5’-ATGGGGAAGGTGAAGGTCG), GAPDH -RV (3’-TAAAAGCAGCCCTGGTGACC), Sox2-FW (5’-AACCAGCGCATGGACAGTTAC), Sox2 - RV (3’-TGGTCCTGCATCATGCTGTAG), Oct4-FW (5’-AGACCATCTGCCGCTTTGAG), Oct4-RV (3’-), NANOG -FW (5’-GGCTCTGTTTTGCTATATCCCCTAA) and NANOG -RV (3’-CATTACGATGCAGCAAATACAAGA).

### Flow cytometry

Cells were collected at the time of plating, and at the 48-hour time-point for each experiment, as well as at the 24-hour time-point for a subset of experiments. All flow cytometry analyses were performed on a BD LSRFortessa™ cell analyser (Becton-Dickinson, San Jose, CA). Cells were dissociated using TrypLE Express (Life Technologies), fixed in the dark for 7 minutes at room temperature in 4% paraformaldehyde, blocked for two hours in 5% FBS and were permeabilised for intracellular staining in ice-cold 70% methanol. All antibodies used were directly conjugated with fluorochrome. Species and fluorochrome-matched isotype controls were used for each antibody to control for non-specific binding.

Cells were incubated with primary antibodies for 30 minutes at room temperature, at the following dilutions; Phycoerythrin-conjugated mouse anti-TRA 1-81 (Ebioscience 12-8883-80), 1:300; Phycoerythrin-conjugated mouse isotype control (Santa-Cruz, SC2870) 1:75; Alexa 488-conjugated Mouse anti-Nanog (BD Bioscience, BD560791), 1:10; Alexa 488-conjugated mouse isotype control (BD Bioscience, BD557702) 1:100; Alexa 647-conjugated mouse anti-Sox2 (BD bioscience, BD56139) 1:80; Alexa 647-conjugated mouse isotype control (BD Bioscience, BD557714) 1:80. Flow cytometry data was collected and preliminarily processed in FACSDIVA™ software followed by analysis and figure-preparation in FlowJo (both Becton-Dickinson, San Jose, CA).

### Proteomics sample preparation

Five biological repeats each consisting of three technical replicates of Rebl.PAT cells were performed, and all analysed together by label-free mass-spec **(Fig. 2A)**. Man-13 samples were analysed by LC-MS/MS in two batches of two biological repeats (each consisting of three technical replicates) to ensure that any identified biomarkers would be resilient to run variation **(Fig. 2A)**. A subset of the cells was also collected and processed for RNA-Seq analysis at the 48-hour time-point.

Media samples were digested using a modified Filter-assisted Sample Preparation (FASP) method [68] with the following modification: Equal volumes of spent media were concentrated to approximately 50 μL with Microcon - 10 kDa centrifugal filter units (Merck Millipore) at a speed of 14,000 x g. This was then washed and centrifuging three times with the addition of 100 mM phosphate buffer pH 7.4 before the proteins were reconstituted in 50 μL of phosphate buffer. The protein concentration was determined using a Millipore Direct Detect® spectrometer and 50 μg (Man-13) or 25 μg (Rebl.PAT) of protein was added to a fresh 10 kDa filter tube with reduction, alkylation and digestion occurring using the filter tubes. After digestion peptides were collected by centrifugation and the samples were desalted with OLIGO™ R3 reversed-phase media [69] on a microplate system dried to completion and reconstituted just before analysis in 5% acetonitrile and 0.1% formic acid.

### LC-MS/MS

Digested samples were analysed by LC-MS/MS using an UltiMate 3000 Rapid Separation LC (RSLC, Dionex Corporation, Sunnyvale, CA) coupled to an Orbitrap Elite (Thermo Fisher Scientific, Waltham, MA) mass spectrometer. Peptide mixtures were separated using a multistep gradient from 95% A (0.1% FA in water) and 5% B (0.1% FA in acetonitrile) to 7% B at 1 min, 18% B at 35 min, 27% B in 43 min and 60% B at 44 min at 300 nL min-1, using a 75 mm x 250 μm i.d. 1.7 μM CSH C18, analytical column (Waters). Peptides were selected for fragmentation automatically by data dependant analysis.

### Identification and quantification of peptides

The acquired MS data was analysed using Progenesis LC-MS (v4.1, Nonlinear Dynamics). The retention times in each sample were aligned using one LC-MS run as a reference, then the “Automatic Alignment” algorithm was used to create maximal overlay of the two-dimensional feature maps. Where necessary a minimal amount of manual adjustment was employed to increase alignment score to above 80%. Features with charges ≥ +5 were masked and excluded from further analyses, as were features with less than 3 isotope peaks. The resulting peak lists were searched against the SwissProt (release 2016-04) and Trembl (release 2016-04) databases using Mascot v2.5.1, (Matrix Science). Search parameters included a precursor tolerance of 5 ppm and a fragment tolerance of 0.6 Da. Enzyme specificity was set to trypsin and one missed cleavage was allowed. Carbamidomethyl modification of cysteine was set as a fixed modification while methionine oxidation was set to variable. The Mascot results were imported into Progenesis LC-MS for annotation of peptide peaks.

### Proteomics data processing

SwissProt and Trembl IDs were used to align data between independent proteomics experiments. Normalised protein abundance data from Progenesis were log2 transformed to improve the normality of distribution. Statistical significance was calculated using Welch’s Two Sample t-tests (p) and adjusted for multiple comparisons using the Benjamini & Hochberg (q).

Peptide sequences for all proteins were uploaded to the SignalP 5.0 server [20] to identify the proportion of proteins bearing a classical secretion signal-peptide sequence. Non-classical secretion was identified by uploading peptide sequences to the SecretomeP 2.0 server [21]; and the recommended cut-off score of 0.6 was used to quantify the proportion of proteins which were likely to be secreted through non-classical pathways. Protein GO-term associations were obtained by from Uniprot, or by use of WebGestalt’s GO-Slim output [70].

### RNASeq data analysis

BAM files were aligned to the human genome using BowTie [71]. In Linux, BAMs were sorted in the command interface using samtools [72], and the number of read counts per gene was obtained using featureCounts [73] (supplemental script 1). For each cell line, a count matrix table was constructed from all the counted BAMs.

The following analysis was conducted in R [74] and full scripts provided (supplemental scripts 2, 3 and 4). Genes with less than 10 reads in all three replicates were filtered out. Counts were normalised using TMM [75] and normalisation factors for each cell line generated. Subsets of filtered counts and normalisation factors were created to represent samples relating to loss of pluripotency. Differential analysis was performed using DESeq2 [76]. Statistically significant differentially expressed genes are defined using FDR corrected p-value <0.05.

### Western blotting

Medium from a variety of cell lines, including Man-1, Man-7 and H9 cells in addition to MAN-13 and Rebl.PAT, was collected as described **in Fig. 1A**, and 3ml of medium was concentrated to 50-100μl in Microcon - 10 kDa centrifugal filter units (Merck Millipore). Samples were run on 10% Bis-Tris gels (Thermo #NW00100BOX) on a Mini Gel Tank (Thermo #A25977) using MES SDS Running Buffer (Thermo #B0002) alongside broad range markers (11-245KDa, NEB #P7712S). Each lane contained 20 μg of protein, heated in Pierce lane marker reducing buffer (Thermo #39000) at 95°C for 10 minutes. Gels were transferred using the IBlot2 Gel Transfer device (Thermo #IB21001), using iBlot 2 Transfer stacks (nitrocellulose membrane, Thermo #IB23001).Cells were analysed by immunoblotting using the following antibodies: 1/200 Mouse anti-Chromogranin A (Novus Biologicalis, NBP2-44774); 1/100 Rabbit anti-COCH (Abcam, ab170266); 1/500 mouse anti-FGFR1 (R&D systems, MAB658); 1/1000 rabbit anti-Follistatin (Abcam, ab157471); 1/1000 rabbit L-Entactin/NID1 (Abcam, ab133686); 1:1000 rabbit anti-Neuronal Pentaxtrin 2 (Abcam, ab191563); 1/1000 rabbit anti-Neurotensin (Abcam, ab172114); 1/300 rabbit L-Olfactomedin-like 3 (Abcam, ab111712); 1/500 mouse anti-Semaphorin 3A (R&D, MAB1250); 1/500 mouse anti-Secreted Frizzled Related Protein 2 (R&D, MAB6838). Secondary antibody staining was performed either using 1/20,000 IRDye 800CW Goat anti-mouse, or 1/20,000 IRDye 680RD Donkey anti-Rabbit (P/N 925-32210 and 925-68071 respectively, both from LI-COR). These fluorescent secondary antibodies were imaged using the Odyssey CLx imaging system (LI-COR) typically in pairs of rabbit and mouse antibodies together. Image brightness/contrast adjustment and densitometry quantification was all performed in ImageJ (NIH).

### Protein extraction method for Western blotting

Media was concentrated using 10KDa Micron filter spin columns (Millipore), centrifuged at 14,000xg for 40mins at 4C, followed by 2 washes with PBS (Life Tech) and spinning at 14,000xg for 15mins at 4C. Protein was resuspended to a total volume of 200μl with 1x PBS. Protein concentration was determined using a BCA Assay kit (Pierce). Concentrated proteins were mixed and vortexed for 2 mins with 1% Trichloroacetic acid-Isopropanol (TCA-IPA) solution (1:10 ratio) and centrifuged 1500xg for 5mins at 5C. Pellets were washed twice with methanol, centrifuged at 800xg for 2mins and resuspended in 6x SDS-PAGE sample buffer, boiled at 96C 10mins and loaded onto an SDS gel for analysis.

### Teratoma formation

Teratoma generation was by subcutaneous injection into Scid Beige mice of approximately 1 million pelleted cells in Matrigel™ [77]. Animals were killed within 11 weeks of injection; skin biopsies fixed in 4% paraformaldehyde and embedded in paraffin wax for histological analysis. Some sections were stained with antibodies to early endoderm (GATA6-CST 5851 1:1600) neurectoderm (Beta3 tubulin R&D Systems MAB1195 1:200) or smooth muscle actin in mesoderm (alpha SMA R&D Systems MAB1420 1:200) followed by incubation with biotinylated secondary antibodies (Vector Labs biotinylated secondaries, anti-mouse, BP-9200 and anti-rabbit IgG BP-9100) and developed using the ABC peroxidase kit (Vector labs PK-6100). Slides were imaged using the 3D Histec Pannoramic250 slide scanner and assessed using CaseViewer with Histoquant licence.

## Acknowledgements

This work was funded by MRC grant MR/M017344 and an MRC Confidence in Concept (CIC) grant to SJK. We thank Qi Wang for the staining in Figure S5, as well as the Histology Facility for equipment used in this study which was purchased with grants from The University of Manchester Strategic Fund. Special thanks go to Peter Walker & Grace Bako for their help with the Histology.

---

**Figure S1.**
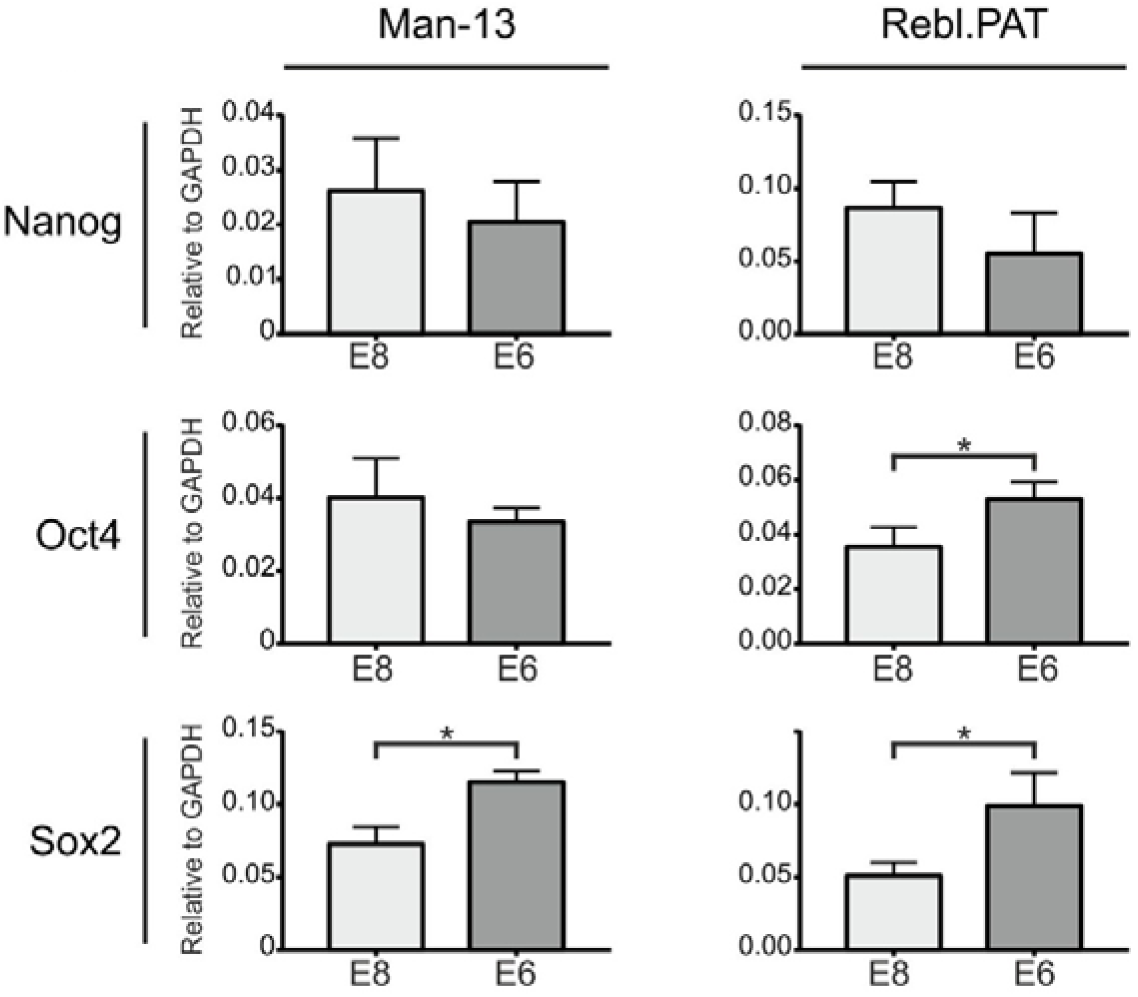
QRTPCR of Man-13 and Rebl.PAT cells cultured in parallel to those used for medium collection, either maintained in E8, or after being incubated for 48 hours in E6. QRT-PCR for pluripotency associated markers Oct4 Sox2 and Nanog relative to GAPDH.

**Figure S2.**
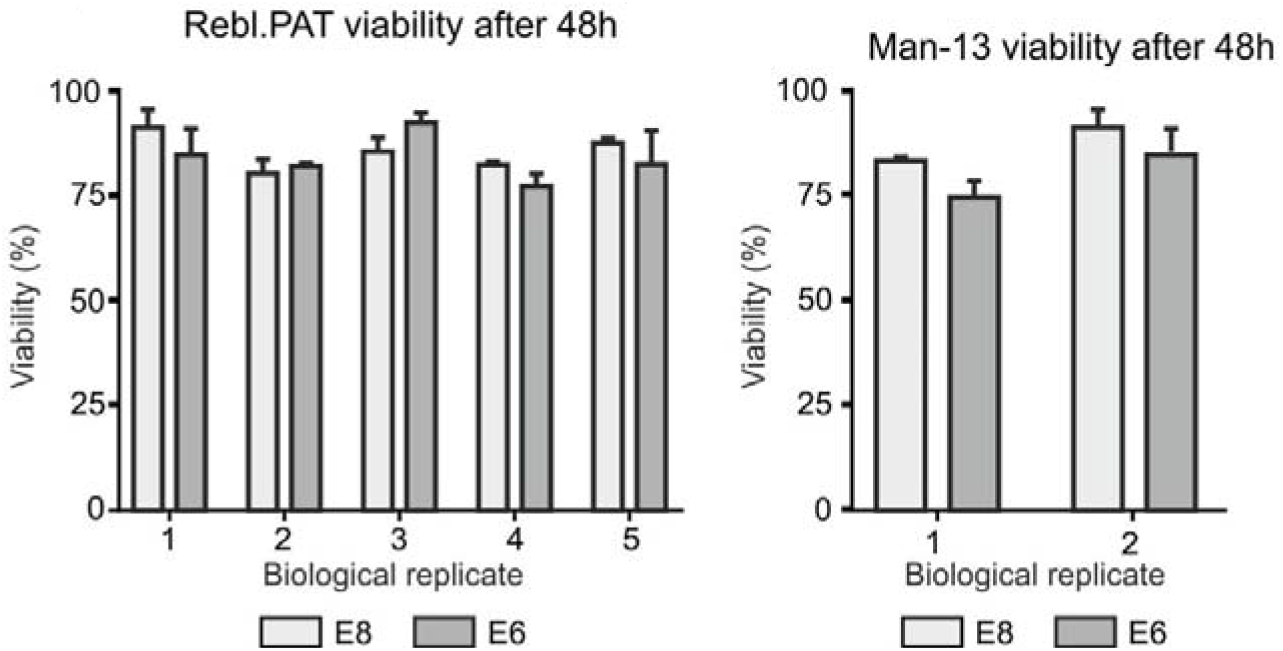
Viability of cells after 48 hours of culture in E6. At the 48-hour timepoint (Fig. 1), viability of E8 and E6 cultured cells was assessed using Via-1 cassettes (Chemometec). All three flasks of cells for each condition were tested, and no significant change between the conditions was observed in any experiment.

**Figure S3.**
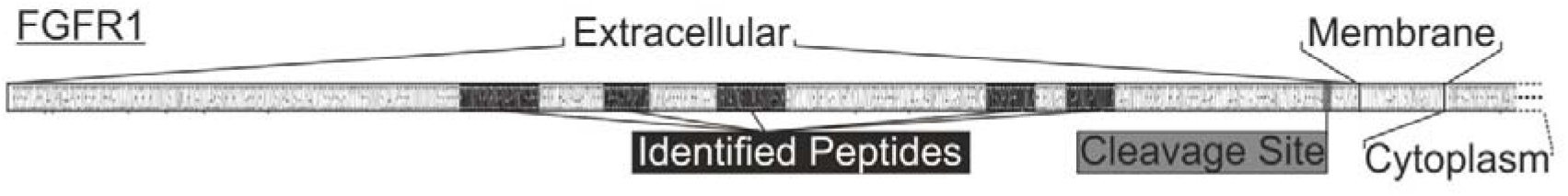
FGFR1 schematic showing peptides used in quantification, and the site of the extracellular cleavage by MMP2 described by Levi, E., et a (1996).

**Figure S4.**
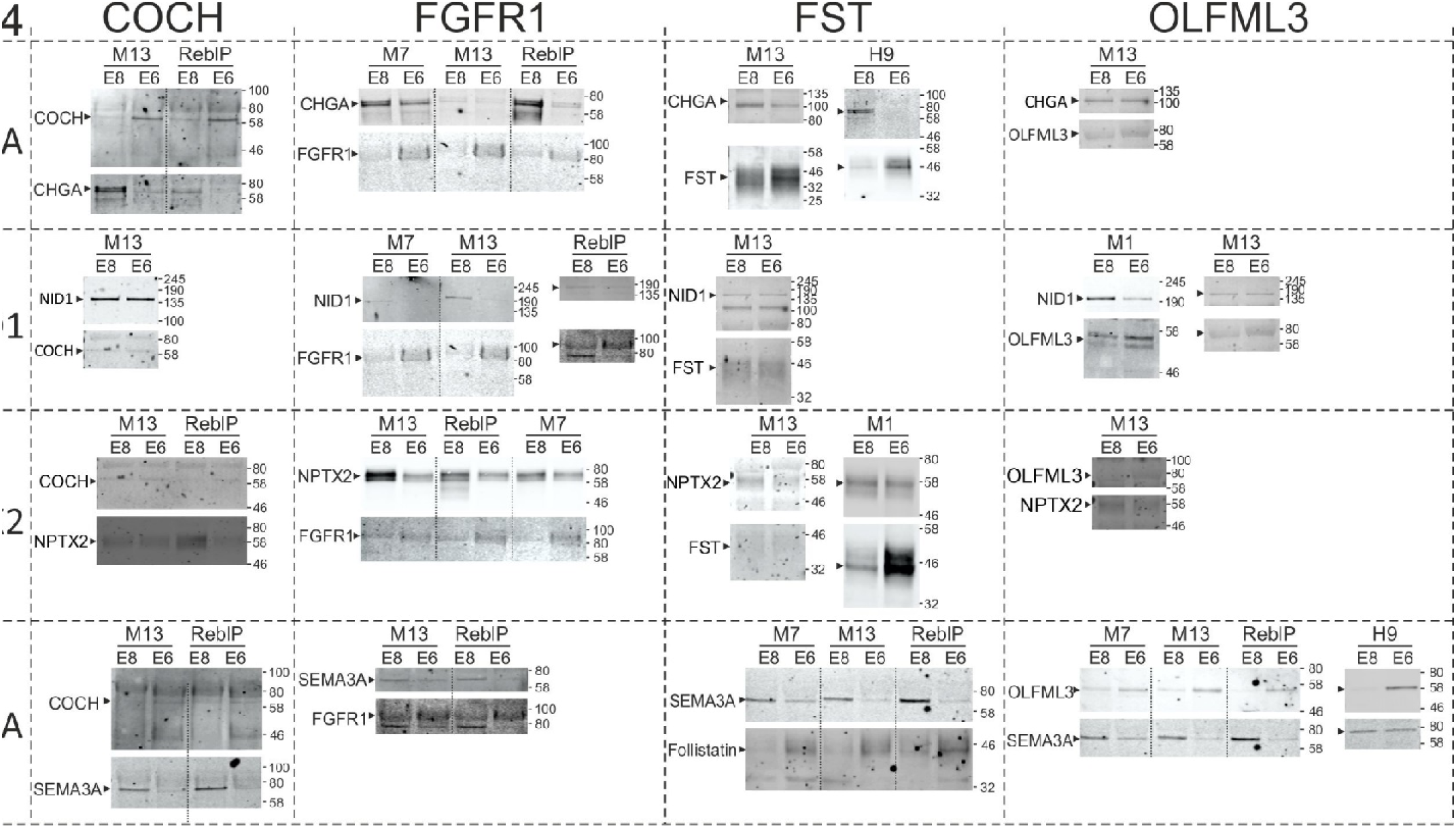
Western blots of marker-proteins in conditioned media. Equal quantities of concentrated conditioned media were run for each condition, and the same membrane probed with both antibodies for each pair. Membranes were imaged using the LICOR Odyssey system. Quantification for these blots is show in **Figure 7A**. E8 enriched proteins on vertical axis E6 enriched proteins on horizontal axis.

**Figure S5.**
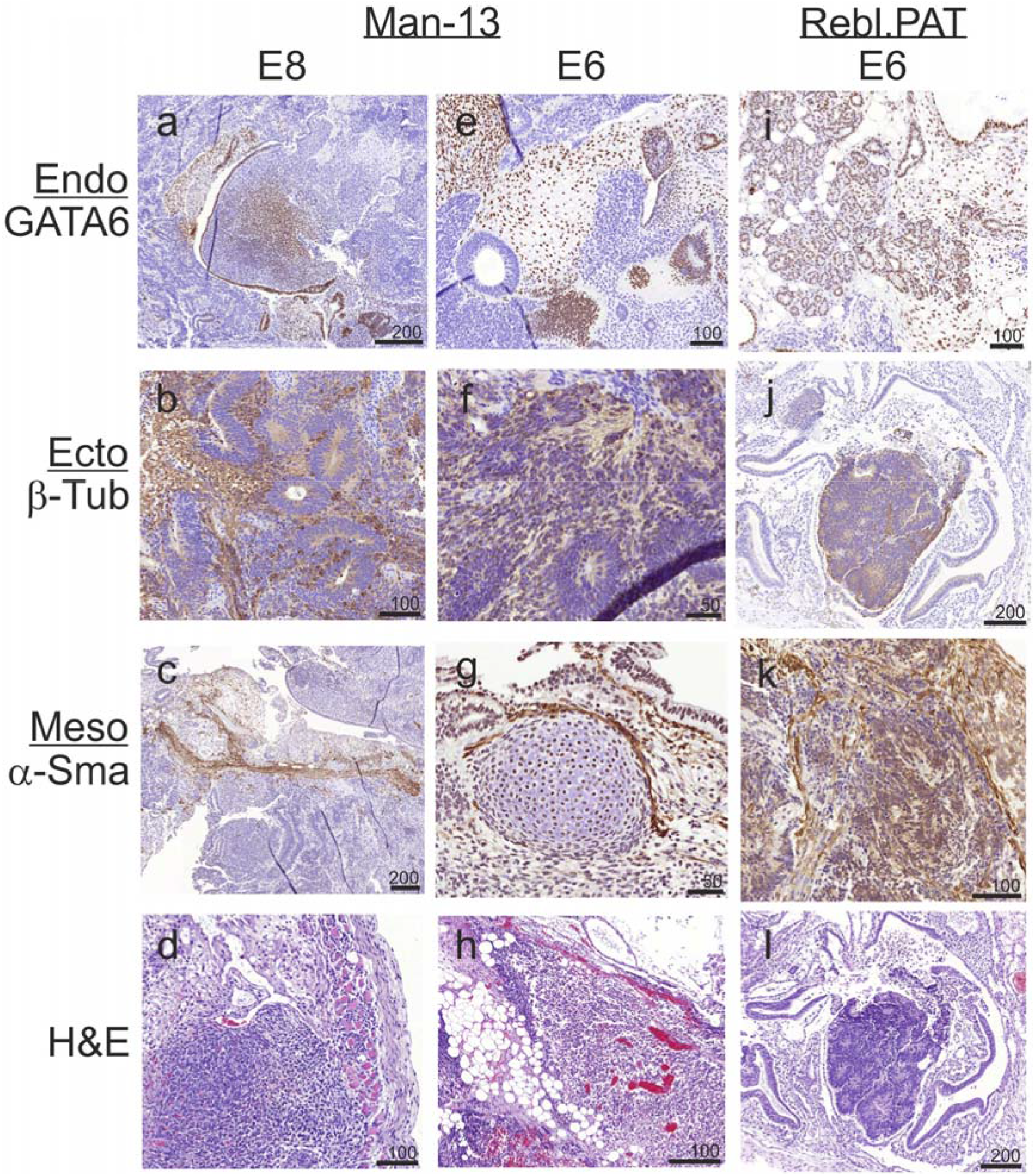
Paraffin sections (7μm) of teratomas generated by subcutaneous implantation of human pluripotent stem cells cultured in in E8 (Man13) or after 2 days in E6 Man13 and Rebl.PAT) stained for morphological and protein markers of the 3 germ layers. Paraffin section were stained with Haematoxalin and eosin or with antibodies to early endoderm (GATA6) neurectoderm (Beta3 tubulin) or smooth muscle actin in mesoderm (alpha SMA) followed by a peroxidase labelled secondary antibody. Scale bar 100um.

**Figure S6.**
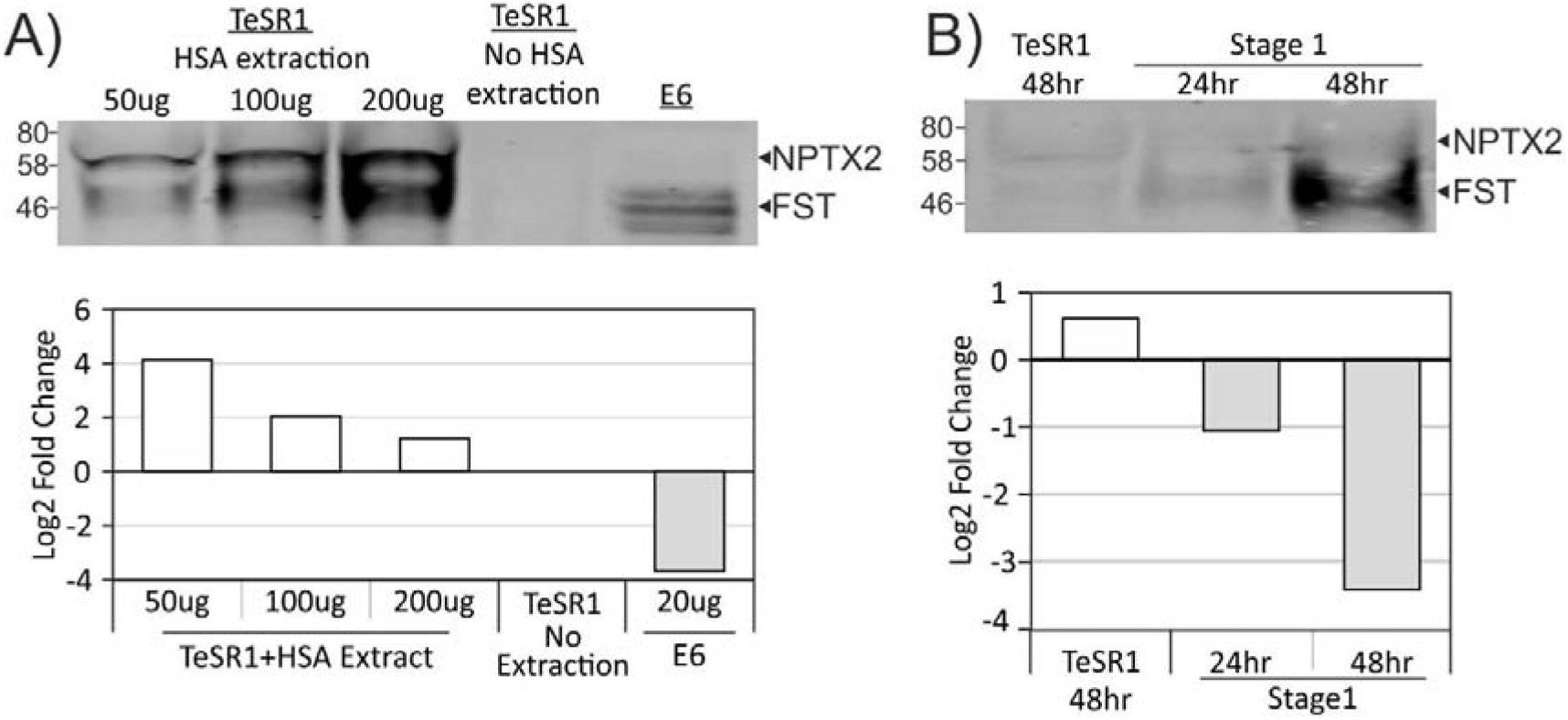
A) NPTX2 / FST Ratio of concentrated TeSR1 pluripotency media from cells grown for 48hrs, and medium subjected to HSA removal using 1%TCA-IPA or without HSA removal. B) NPTX2 / FST Ratio in media concentrated from cells grown in TeSR1 pluripotency medium versus mesodermal (stage1 chondrocyte) differentiation medium showing that the change in protein ratios for NTPTX2 and Follistatin holds in both experiments if HSA is removed.

